# Evolving social behaviour through selection of single-cell adhesion in *Dictyostelium discoideum*

**DOI:** 10.1101/2021.07.15.452515

**Authors:** Sandrine Adiba, Mathieu Forget, Silvia De Monte

## Abstract

The social amoeba *Dictyostelium discoideum* commonly forms chimeric fruiting bodies. Genetic variants that produce a higher proportion of spores are predicted to undercut multicellular organization unless cooperators assort positively. Cell adhesion is considered a primary factor driving such assortment, but evolution of adhesion has not been experimentally connected to changes in social performance. We modified by experimental evolution the efficiency of individual cells in attaching to a surface. Surprisingly, evolution appears to have produced social cooperators irrespective of whether stronger or weaker adhesion was selected. Quantification of reproductive success, cell-cell adhesion and developmental patterns, however, revealed two distinct social behaviours, as captured when the classical metric for social success is generalized by considering clonal spore production. Our work shows that cell mechanical interactions can constrain evolution of development and sociality in chimeras, and that elucidation of proximate mechanisms is necessary in order to understand the ultimate emergence of multicellular organization.

## INTRODUCTION

Questioning about the evolution of unicellular organisms has long focused on properties of individual cells that live in isolation. In the recent decades, it became clear that many functions of microbial populations – spanning from stress resistance in biofilms and invasion by a pathogen to spore production in aggregative microbial species – derive from the collective, social outcome of interactions among cells (Ackermann, 2015). Understanding how selection shapes functionality of groups of cells is challenging, in particular when multicellular aggregates are composed of multiple cellular types (Rainey and De Monte, 2014). In this case, according to a common distinction, ‘cooperator’ and ‘cheater’ types compete within groups and can disrupt their collective functions.

The ‘social amoeba’ *Dictyostelium discoideum*, whose life cycle comprises a multicellular body formed by aggregation of formerly independent cells, is an established model system to address the role of genetic conflicts on the evolution of multicellular organization (Strassmann et al., 2000; Li and Purugganan, 2011). Chimeric aggregates, where cells of different strains develop together and differentiate into several tissues, readily occur both in nature and in the lab (Fortunato et al., 2003b; Gilbert et al., 2007; Sathe et al., 2010; Castillo et al., 2011). Within aggregates, a cell’s terminal differentiation has two main issues: spores, that disperse and survive starvation, and a stalk of dead cells. The dramatic disparity in reproductive output of such two cellular fates generates strong conflicts within chimeric aggregates. Consistent with theoretical expectations, “cheater” strains that are over-represented in the spores have been shown to get enriched in successive cycles of aggregation and dispersal (Strassmann et al., 2000; Gilbert et al., 2007; Santorelli et al., 2008). Assortment of cells of similar social investment is believed to be the essential factor in curtailing the long-term evolutionary success of such “cheaters”, as it prevents them to reap collective benefits (Strassmann et al., 2000). In practice, several mechanisms can produce such assortment (e.g. spatial structure, differential adhesion, kin recognition) (Strassmann and Queller, 2011). It is still unclear however what cell-level features implement in *D. discoideum* the general organizing principles that underpin the repeated emergence of aggregative multicellularity (Grosberg and Strathmann, 2007; Márquez-Zacarías et al., 2021; Forget et al., 2021).

Cell adhesion has been invoked as a primary means to achieve assortment among different strains (Hirose et al., 2011; Queller et al., 2003; Ispolatov et al., 2012; Garcia and De Monte, 2013; Garcia et al., 2014, 2015; van Gestel and Wagner, 2021). Beside being ubiquitous in the microbial world and essential for cell sorting in multicellular organisms, it is involved in every step of *Dictyostelium* life cycle. Amoebae spend most of their time in a vegetative phase, where isolated cells crawl, divide, and feed individually on bacteria (Kessin, 2001; Raper, 2014). Adhesion to their physical support determines their motility properties and changes throughout the cell cycle (mitotic cells detach from the surface) (Nagasaki et al., 2002; Plak et al., 2014). During the multicellular development, that is triggered by nutrient depletion, different forms of cell-cell adhesion come to the fore. At the beginning of aggregation, cells display non-polar cell-cell adhesion. Later, they attach head-to-tail while the streams they form converge towards the so-called mounds (Gerisch, 1980; Fujimori et al., 2019). These develop in a matrix-coated slug that displaces by collective cell motion, and finally forms a fruiting body. Despite not being independent (Wang et al., 2014), cell-substratum and cell-cell adhesion are associated to different gene expression profiles (Rosengarten et al., 2015). Importantly, by their implication in different stages of the life cycle, different forms of adhesion are subjected to distinct selective pressures. Selection of cell-cell adhesion presupposes a multicellular context, whose collective function will generally depend on the type and proportion of its constituent cells. Cell-substratum adhesion, on the other hand, is primarily influenced by the physical environment that cells experience, and is expected to evolve independently of the quality and (to a certain extent at least) quantity of other cells present in the surroundings.

With this distinction in mind, we examined whether the social behaviour of *Dictyostelium* would be affected by selection acting on adhesion of single cells to a surface. We therefore designed a directed evolution experiment where vegetative cells were selected based on their adherence to a culture vial. Contrary to trait modifications obtained by mutagenesis and KO experiments, artificial selection allows to mimic gradual phenotypic changes that are expected to occur in nature, and that may involve multiple genes simultaneously, as well as epigenetic changes. Moreover, it allows to track the evolutionary history of adaptations. This study thus connects proximate mechanisms of evolved behaviours – such as cell adhesion and dynamics during development – to their consequences for the reproductive success in chimeras, which underpin ultimate causation of the evolution of social behaviour.

Starting from the same ancestral AX3 strain, we selected cells for increased or reduced adhesion to a surface during two months of daily passages. Three replicate lines were evolved independently under opposite selective protocols and acquired heritable, consistent within treatments, variations in adhesion. In binary chimeras composed of the evolved lines with their common ancestor, the derived lines always produced a smaller fraction of spores, irrespective of their being more or less adhesive to the surface. In order to understand the reason of such surprising evolution of cooperative behaviour through selection on individual cell properties, we characterized the phenotype of the evolved lines both in isolation and during development. These observations allowed us to interpret the developmental dynamics in chimeras and to identify two opposite routes by which evolved lines achieve a same, apparently cooperative, behaviour. In order to distinguish between these two routes, we propose a metric of social performance, alternative to simple bias in spore frequency, that accounts for the observed developmental differences. We conclude discussing the potential implications of selection for single-cell mechanical properties on the evolution of social behaviour.

## RESULTS

*D. discoideum* cells were subjected to a directed evolution experiment, where selection was applied to their adhesion to a surface (see Methods: Evolution of cell-substratum adhesion). Serial passages to fresh medium were realized by separating cells that remained in suspension from cells attached to the bottom of the culture flask. The first and the second were selected in the Top and Bottom treatment, respectively. Nutrients availability was sufficiently large to keep cells in the exponential phase of growth, thus in their vegetative unicellular state. After 30 rounds of selection (corresponding to 85-97 generations, see Figure S1), we obtained three lines for each of the two treatments, that in the following we refer to as lines 1, 2, 3 and Top and Bottom. We expect these derived lines to harbour genetic or epigenetic variations that underpin differences in adhesion with respect to the Ancestor AX3 strain. This was confirmed by the observation that phenotypes were reproducible even after multiple cycles of growth and dilution. In order to minimize differences in physiological state, all the experiments on a same cell line were realized from the same frozen stock after thawing, growth and dilution to a same cell density. Population-level assays described in the following were realized for the three independent evolved lines of the Top and Bottom treatment, in three biological replicates started from the same initial culture, while time-lapse movies were captured in duplicates.

### Cell-substratum adhesion and social behaviour of the evolved lines

As expected, at the end of the evolution experiment, Top, Bottom and Ancestral cells differed in their adhesion to the surface, assessed in early exponential growth phase (see Methods: Cell-substratum adhesion assay). The three evolved Top lines were twice less adhesive than the Ancestor, while the three evolved Bottom ones displayed a 20% increase in adhesion (Figure 1 A). Within each treatment (Top and Bottom) we found no significant difference in cell-substratum adhesion (Student *t-test*).

**Figure 1.**
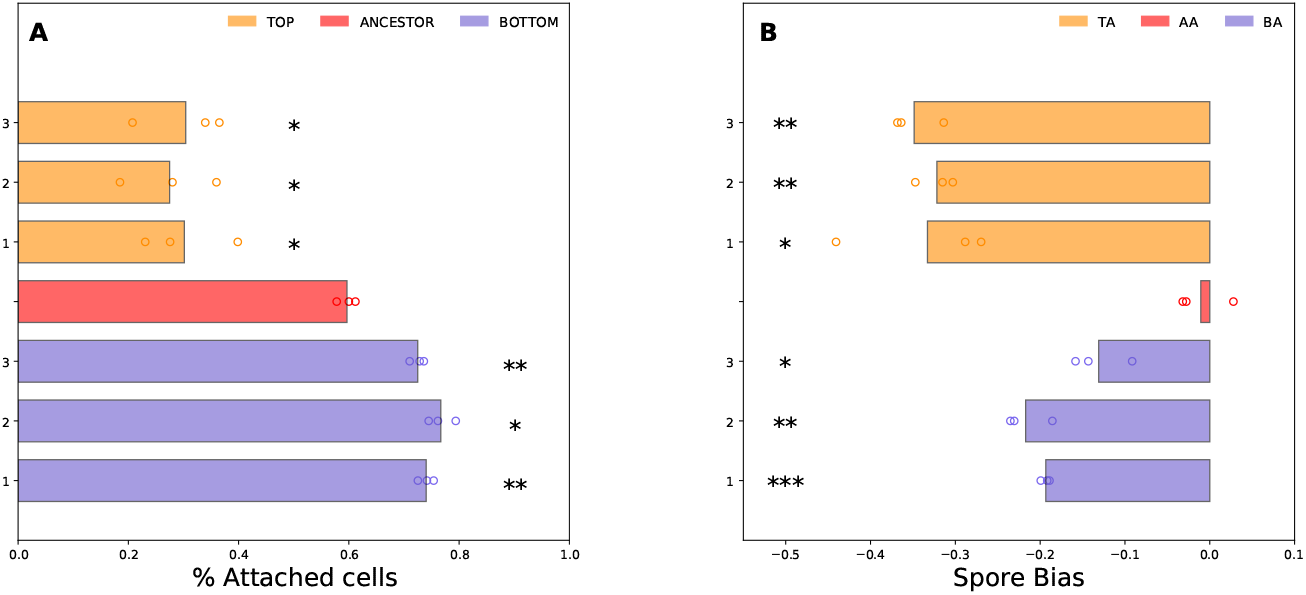
Cell-substratum adhesion and spore bias for three evolved Top and Bottom lines. (**A**) Cell-substratum adhesion for three evolved Top and Bottom lines. Top (orange) and Bottom (violet) lines were less adhesive and more adhesive, respectively, compared to the Ancestor (red). (**B**) Bias in spore allocation of the focal line relative to its initial proportion of cells in the mix (as estimated from Equation 2). Chimeras were formed by mixing RFP-tagged with GFP-tagged Ancestor cells (AA: red), and RFP-tagged Ancestor with the three Top lines (TA: orange), and the three Bottom lines (BA: violet). Top and Bottom contribute significantly less than the Ancestor to the spore pool. Circles represent data points of three biological replicates of the same chimeric mix, Student *t-test* : * *p<*0.05, ** *p<*0.001, *** *p<*0.0001.

We then wondered whether the selected differences in substratum adhesion of single vegetative cells would also result in different social behaviour. Classically, social behaviour is assessed in chimeras by measuring the proportion of spores produced by each of the two strains. Deviation with respect to the composition of the initial mix (typically 50% of cells of each strain, Equation 2 in Methods: Measures of spore formation efficiency and metrics of social performance)(Fortunato et al., 2003a; Foster et al., 2004; Santorelli et al., 2008, 2013) determines how much one type is enriched in the spores.

We mixed evolved Top and Bottom lines and the Ancestor strain (all marked with GFP) with the Ancestor strain marked with RFP. In the following, we refer to chimeras where Ancestor cells were mixed to an equal amount of Bottom (Top, Ancestor) cells as BA (TA,AA, and lines 1, 2, 3, respectively). We quantified the composition of the spore pool by counting the total number of spores and measuring the fraction of spores belonging to each type (see Methods: Measures of spore formation efficiency and metrics of social performance). First, we used the AA mix to check that differences in fluorescent labelling plasmid did not per se induce a spore bias (Figure 1 B).

When mixed with the Ancestor strain, all evolved lines contribute to the spore pool less that their initial proportion in the mix (Figure 1 B). The three Top lines and three Bottom lines scored consistently against the Ancestor, with Top lines displaying a stronger reduction in reproductive success with respect to the Ancestor. Based on the widely used social metric Equation 2, all the evolved lines thus behave as ‘cooperators’ towards the Ancestor. Indeed, they are under-represented in the spore pool compared to the latter, hence they provide a relative fitness advantage to the Ancestor. Higher levels of sociality may have been expected in cells that adhere more to a surface if they also adhere more efficiently to one another, as was also previously observed in *Myxococcus* (Velicer and Yuen-Tsu, 2003). It is more puzzling, instead, that a similar result is also observed for the less adhesive Top lines. However, chimeras with both Top and Bottom lines produce significantly less spores than the Ancestor (Figure S2), indicating that, while in relative terms the Ancestor is always advantaged, the evolved variations in adhesion have a deleterious effect on the total reproductive output.

In the following, we look for the self-organization mechanism at play when similar social behaviour is produced, during development, by cells with opposite adhesion to the surface.

### Cell adhesion to a surface affects multicellular development

The multicellular stage of the life cycle was not directly subjected to selection. In the absence of pleiotropic effects of cell-substratum adhesion, its traits should therefore be chiefly affected by drift, thus they should not undergo directional evolutionary variation. In this section, we investigate the developmental life cycle of the derived lines and show that it differs with respect to their common ancestor via related variations in cell-cell adhesion.

As a first step, we observed clonal multicellular development for the three independently evolved lines of the Top and Bottom treatments. All experiments were started from the same initial cell density, equal to the total density in chimeras (see Methods: Developmental life cycle). Derived lines are still able to perform the developmental life cycle from aggregation to the formation of the fruiting bodies (Movie S3: Clonal aggregation of the Ancestor, Top (line 1) and Bottom (line 1)), except the Top line 3 whose development stops at the mound stage. However the timing of each stage (aggregation, streaming, mound, tipped aggregate, slug and Mexican hat) varies considerably between Top and Bottom treatments and is consistent among the three independently evolved lines of a same treatment (Figure 2).

**Figure 2.**
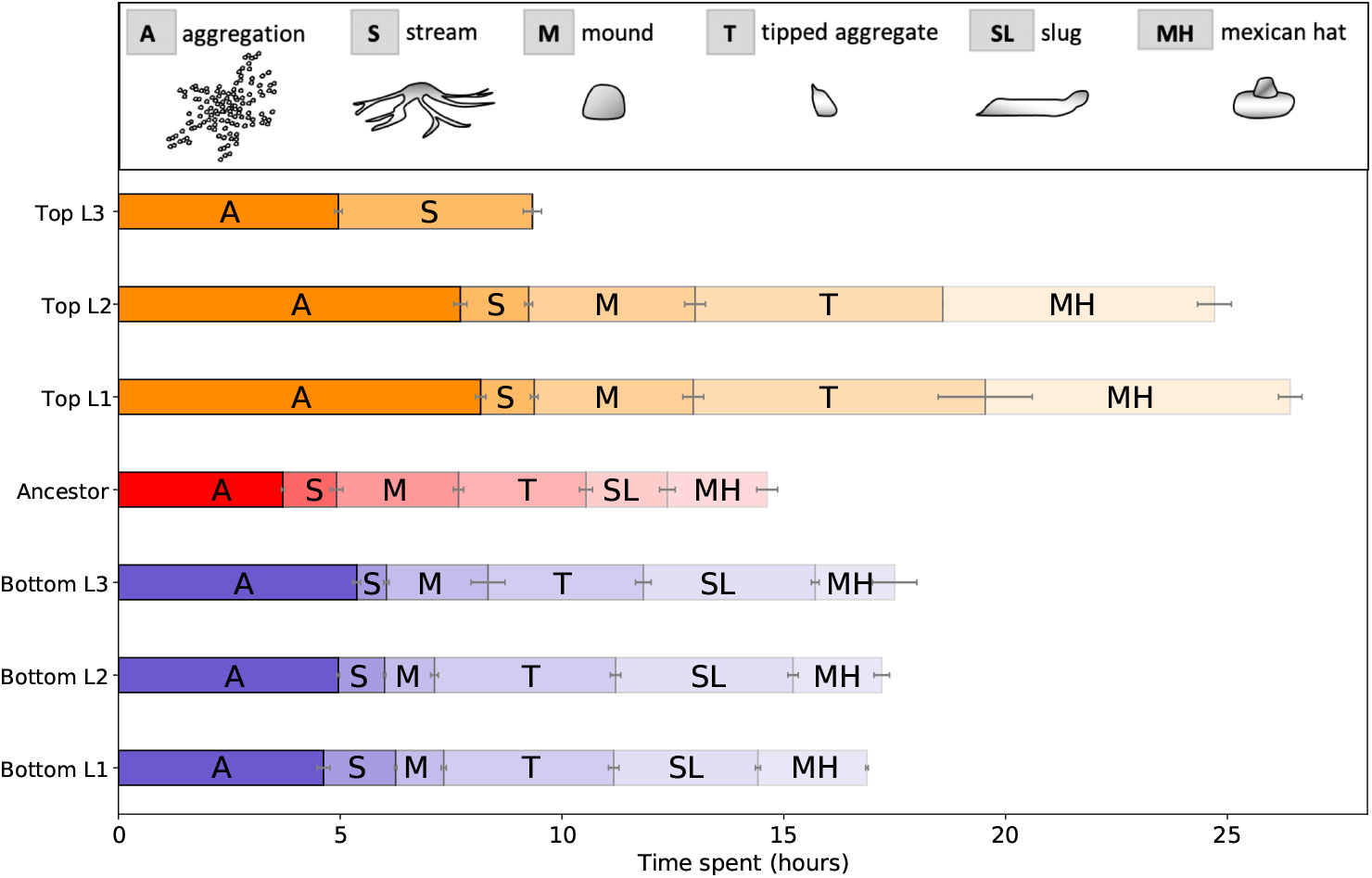
Timing of developmental life cycle. The different stages of the developmental cycle were classified by analysing phase contrast time-lapse movies (see Methods: Time-lapse microscopy and image analysis). The figure displays the mean of two independent experiments for the Ancestor and each Top and Bottom line. Differences in development, most notable for the Top lines, are discussed in the text. Colour code as in Figure 1.

Figure 3 shows snapshots of such different stages in the Ancestor, and for representative Top and Bottom line 1 (see Figure S4 A for snapshot of evolved lines 2 and 3). The developmental cycle of Bottom cells, that were more adherent to the surface in exponential phase, is similar to that of the Ancestor strain. Five hours after plating, the firsts streams are observed. For the same initial cell density, the Bottom lines tend to form a smaller number of aggregates (Figure S5 B). Mounds have, on average, a smaller diameter (Figure S5 A). This does not necessarily mean they contain less cells, as differences in the area covered by aggregates may be due to the influence of cell-substratum adhesion on the shape of the mound. Slugs of the Bottom lines migrate longer and fruiting bodies are noticeably taller, indicating a higher propensity of Bottom cells to differentiate in the stalk.

**Figure 3.**
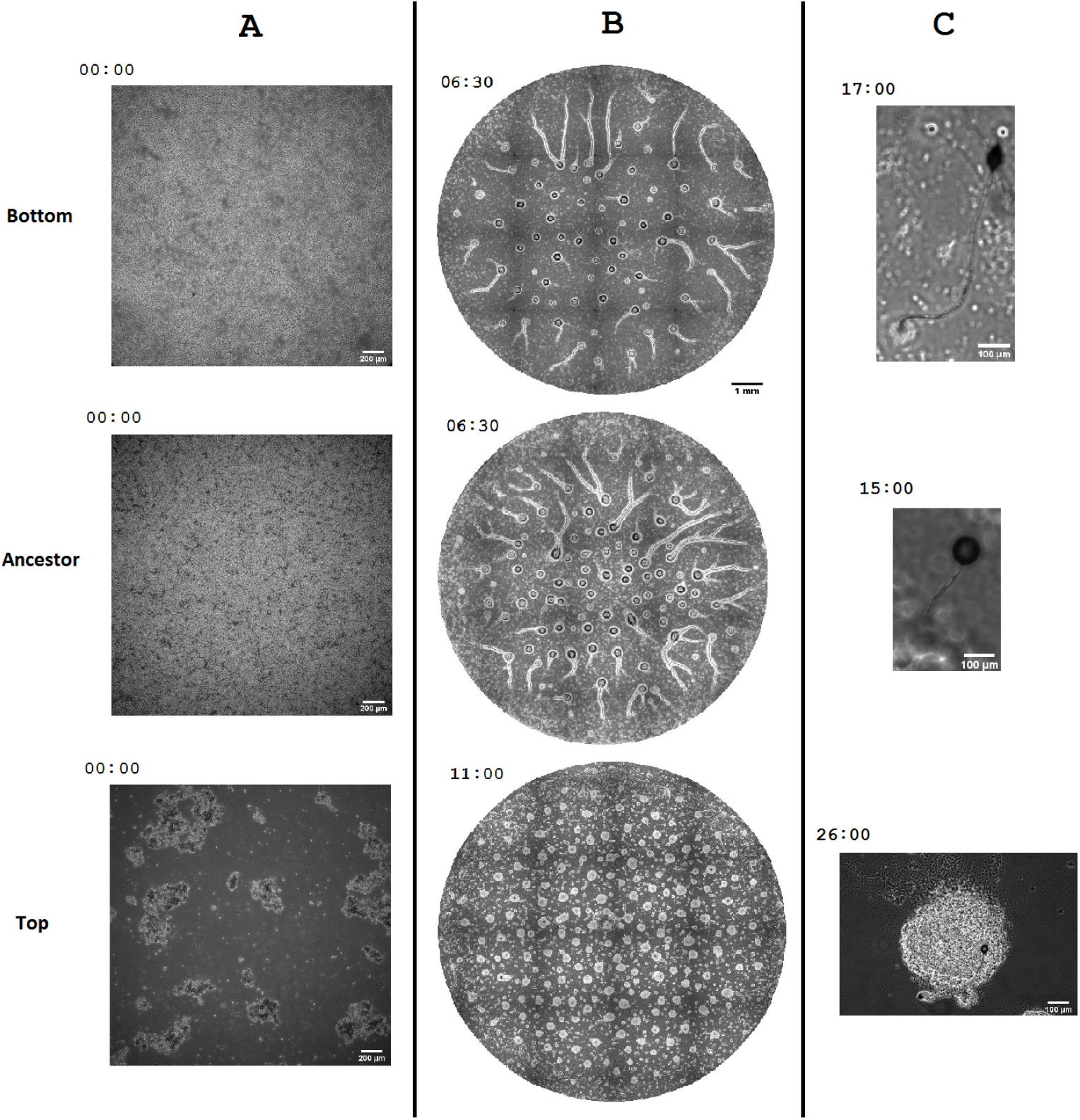
Developmental cycle of the Top, Ancestor and Bottom line 1. Phase contrast images of three stages of the cycle: beginning of aggregation (**A**), mounds (**B**), fruiting bodies (**C**). Snapshots are taken at equivalent developmental phases, rather than at a same absolute time, in order to meaningfully compare the different treatments (see Methods: Time-lapse microscopy and image analysis). Pictures represent line 1 of the Top and Bottom treatments and are representative of the behaviour of all lines of the same treatment (except for the Top line 3 whose life cycle stopped at the mound stage without fruiting body formation, see Figure S4 A for snapshot of evolved lines 2 and 3). The time of capture is indicated on the left of each picture (see Figure 2 for reference). As discussed in the text, lines selected for their adhesion to the surface differ significantly from the Ancestor also in their developmental cycle, with the Top lines displaying the largest variations.

The Top lines, on the other hand, is more severely impaired in its development (Figure 3). From the beginning, Top cells tend to float and readily form small clusters (Figure S5 A). Streaming starts 7.5 hours after plating, much later than for the Ancestor. Similarly, the first stages of multicellular development are considerably delayed. Streams tend to break, mounds are more numerous and of smaller area (Figure S5 A-B). The most notable qualitative difference with respect to the other treatment and the Ancestor is that there is no slug stage, and tipped mounds directly develop into small fruiting bodies, leaving off numerous cells in the proximity of the basal disc. Moreover, formation of the fruiting body often fails, suggesting that reduced cell-substratum adhesion plays a key role in the final stages of multicellular development. Particularly, it appears to impair the organization of the tip, which defines the developmental axis of the slug in the Ancestor (Rubin and Robertson, 1975), and the anchoring of the sorocarp onto the surface. In the Top line 3, moreover, multicellular development does not proceed beyond the mound stage.

Altered developmental patterns in the evolved lines resulted in a lower number of spores produced (Figure S6). The observed differences can be attributed either – for Bottom cells – to the formation of fruiting bodies with a disproportionate fraction of stalk cells or – for Top cells – to frequent failure to complete multicellular development, which we expect to be at least partially rescued in chimeras. Cell-substratum adhesion thus affects non-monotonously reproductive output in genetically homogeneous multicellular aggregates.

### Differences in cell adhesion to a surface affect cell-cell adhesion during development

Modifications of cell-substratum adhesion in the selected lines can influence multicellular development either directly, or indirectly through some co-varying trait that has direct developmental effect.

We first investigated how cell-surface adhesion properties varied during the multicellular, ‘social’ cycle. The analysis was realized for the line 1 of the Top and Bottom treatments as detailed in the SI (Figure S7). Cell-surface adhesion was measured at the onset of aggregation (‘Starvation’) and at the time when streams started to form (‘Streaming’) (Figure S7). Compared to the previously characterized vegetative phase, adhesion strength increases first and then decreases at the later developmental phase. Despite this temporal modulation, adhesion differences between Top and Bottom cells remained consistent.

Although adhesion to a substratum could directly affect cell positioning during development, it is also possible that multicellular development gets steered through pleiotropic, indirect effects on cell-cell adhesion (Wang et al., 2014). The latter were assessed by quantifying the capacity of single cells in suspension – thus independent of their substratum adhesion – to form clusters (Gerisch, 1968) (see Methods: Cell-cell adhesion assay).

We observed that cell-cell adhesion was consistently lower in the Ancestor than in the derived lines at starvation (Figure S8). However, increased cell-cell adhesion appears to rely on different mechanisms for Top and Bottom cells (for details see Figure S8). Bottom cells are more adhesive at later developmental stages, which involve proteins that are responsible for polar contacts. Thus, they maintain a polar adhesion similar to the Ancestor during the streaming stage. On the other hand, Top cells are more adhesive during the early development, while they are likely to develop weaker polar contacts.

These observations support the notion that adhesion differences are a primary cause of the previously discussed developmental variations in clonal multicellular aggregates.

### Developmental patterns of evolved lines in chimeras and social success

Understanding how selection acting on the unicellular phase of the life cycle affects social behaviour, thus how derived lines end up being under-represented in the spore pool, requires examining the developmental cycle of chimeras. Evolved lines are marked with GFP. This allows us to observe how cells of different treatments behave during development in mixes with an equal quantity of Ancestor cells, that we marked with RFP. For all binary mixes, cells were deposited on phytagel so that labelled cells were homogeneously distributed at the beginning of development. The developmental cycle was then followed by time-lapse microscopy for all evolved lines.

Anomalies in developmental timing observed in pure Bottom and Top cultures were largely rescued by mixing with the Ancestor (Figure S10), as often happens with mutants deficient in some multicellular function. Most notably, Top cells, that do not form slugs on their own, recovered the normal succession of developmental phases, though their development was still slightly slowed down.

### Differential adhesion affects cell aggregation and sorting during streaming and within slugs

In order to compare the behaviour of Top and Bottom lines in chimeras with the Ancestor, in the following we describe how Top and Bottom lines behave in successive stages of the chimeric developmental life cycle. We take again line 1 as representative of each treatment (see Figure S4 B for snapshot of evolved lines 2 and 3). Looking at the position of cells, we establish connections with previously characterized differences – among treatments and during developmental time – in cell-substratum and cell-cell adhesion, and elucidate the mechanistic bases of the observed social behavior (Figure 4).

**Figure 4.**
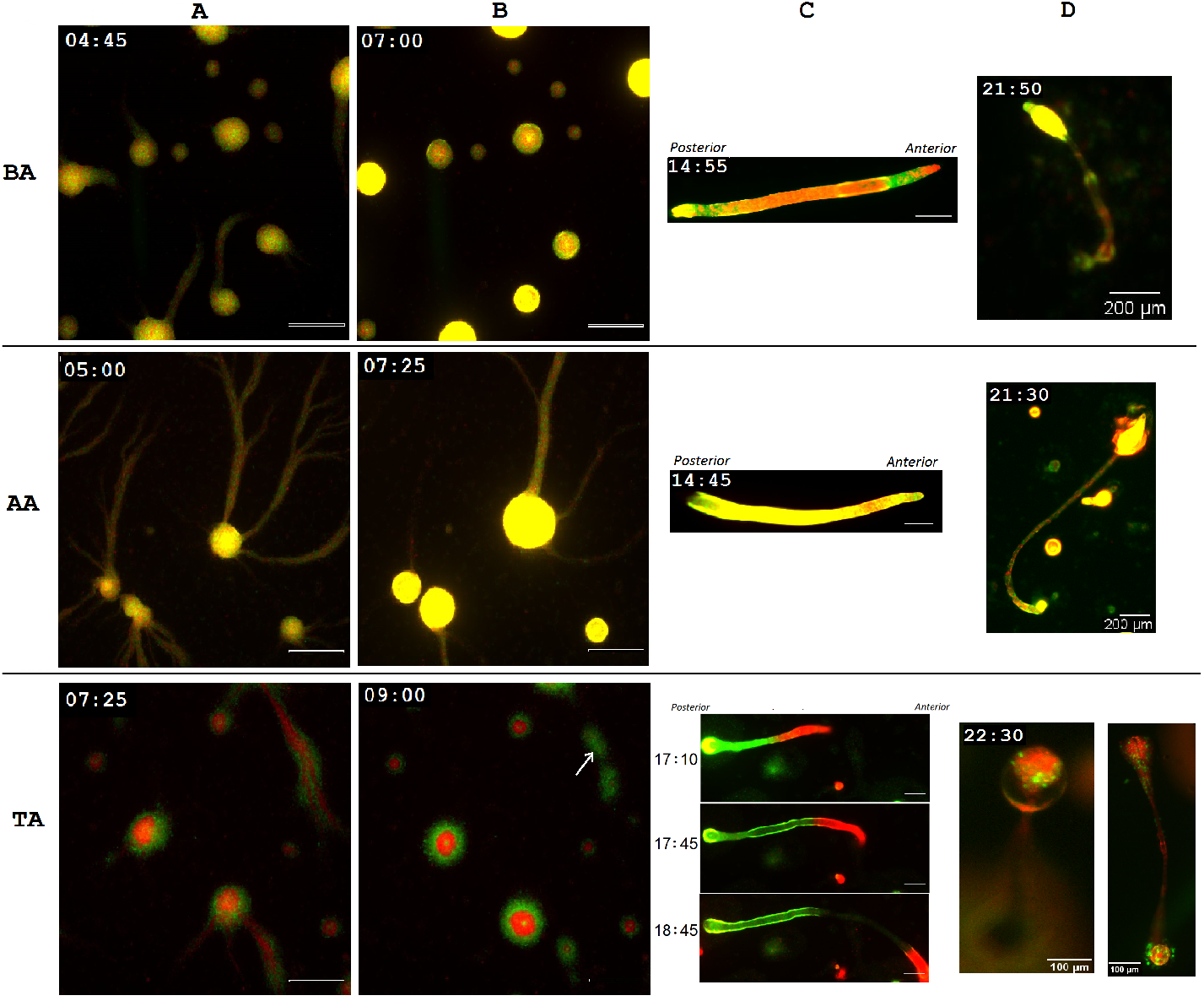
Developmental patterns in chimeras. Snapshots from time-lapse movies during multicellular development in chimeras BA (top), AA (middle) and TA (bottom) for evolved line 1 which is similar to other evolved lines (2 and 3, see Figure S4 B). Ancestor strain is marked with RFP (and GFP for control) and evolved lines with GFP. Cultures with an equal proportion of each type were starved and plated at the same total density (Methods ‘Developmental life cycle’). (**A**) In Early aggregation, Top cells are positioned at the edge of the streams and at the periphery of aggregates (Scale bar = 500 μm). (**B**) Cells that enter the mound last (the arrow indicates a stream in the late aggregation stage) are predominantly Top cells. In the mound, Top cells are found in the center and periphery of the aggregate (Scale bar = 500 μm), producing a bull’s eye pattern. (**C**) In the slug, Bottom cells are mainly localized in regions that later develop into upper cup and basal disk, while Top cells are enriched at the back of the slug. The slug regularly breaks off at the point of transition between Top-enriched and Ancestor-enriched regions. The separation typically happens just after the slug emerges from the mound, so that a significant fraction of Top cells remain in proximity of aggregation centers, but they do not develop further. Scale bar = 200 μm. (**D**) In the fruiting bodies, Bottom cells are concentrated in the tip, the upper cup, and the basal disk, and Top cells are prominently found in the spore ball, and in the basal disc.

BA chimeras display no bias in cell localization before the slug stage. In TA mixes, Top cells are able to enter streams, but they localize predominantly to their exterior, while traction appears to be mostly provided by Ancestor cells that attach more efficiently to the surface. Even though Top cells appear to join aggregates concomitantly with Ancestor cells, streams that reach the mound latest are almost exclusively composed of Top cells. This physical segregation, likely the result of passive physical sorting, is also evident in the mound phase. During development of the Ancestor, cells – previously organized in head-tail chains inside the streams – turn around the aggregate center, maintaining their polar orientations. Top cells, that have low polar adhesion, get excluded from the rotating ring and accumulate at the outer boundary of the mound and in its center. This gives to the mound a bull-eye appearance (Figure 4 B (TA)).

Bottom and Ancestor cells appear interspersed in BA chimeric mounds, reflecting the capacity of both strains to establish polar contacts (though with different intensity). However, differences in surface adhesion and, possibly, in cell-cell adhesion affect cell sorting within slugs. Bottom cells tend to be enriched in the pre-stalk, anterior region of the slug (but not on the very tip) and in the posterior zone (but not the very rear). Similar to pstAO and pstAB cells, that compose the back part of the slug head in normal development, they appear to migrate towards the rear of the slug and eventually contribute disproportionately to the basal disc and the upper cup. Both these positions require strong adhesion: the basal disk keeps the fruiting body attached to the surface, and the upper cup ‘pushes’ the spore mass along the stalk. Bottom cells thus behave as expected for ‘cooperators’, as their adhesion properties make them take on roles that are fundamental for the structural stability and function of the sorocarp.

In TA chimeras, the very distinctive bull-eye pattern of the mound reflects in the later geometry of the slug. From the tipped aggregate stage, it is the Ancestor cells that drive the formation of the tip and the emergence of the slug, whereas Top cells tend to remain to the margins. Even if they have weaker polar attachment, they are carried along by Ancestor cells, whose intact cell-substratum adhesion propels the collective motion of the multicellular stage. Top cells are thus eventually found in the back part of the slug, that is in the pre-spore region, thus providing them a potential advantage over the Ancestor strain. However, a sizeable posterior portion of chimeric slugs regularly breaks off at the point of transition between Top-enriched and Ancestor-enriched regions (Figure 4, TA, C.).

The detached fragments, that lost the traction provided by Ancestor cells, and possibly also the capacity for efficient sorting, do not develop any further. Top cells that remain in the slug end up almost exclusively in the spore mass. At the population level, this is nonetheless insufficient to compensate for the earlier segregation. So, less adhesive cells behave, within single slugs, like extreme cheaters, as the collective functionalities are entirely provided by Ancestor cells. On the other hand, differential adhesion and the subsequent spatial sorting protects Ancestors from the extremely asocial behaviour of cheaters. Top cells behave analogously to the *csA* knockout mutant, whose reduced attachment to other cells has been interpreted as a ‘green beard’ signal. Similar to cells evolved to be less adhesive to the substrate, the *csA* knockout mutant is found in excess among the spores, but it is largely excluded from cooperative streaming when mixed with wild type cells (Queller et al., 2003). It is to be expected that the efficiency of segregation depends on several factors other than differential adhesion, most importantly the fraction of Top cells in the population. When they are in small proportion, indeed, it is possible that Top cells manage to stick to the slug, thus obtaining a disproportionate advantage over the Ancestor. Further investigations of this hypothesis are under way.

Observing their development in chimeras, we saw that Bottom and Top lines differ profoundly in the way they become underrepresented in the spores. Bottom cells behave as one would expect: their increased adhesiveness makes them more likely to bear the cost of sociality by disproportionately contributing to the structural stability of the fruiting body. Top cells, on the other hand, impair both the Ancestor’s and their own development, but they end up bearing the larger share of damages. In the last paragraph, we will consider a metric that allows to differentiate these two behaviours.

### Spore Formation Efficiency and deviation in spore allocation in chimeras

The qualitative differences in spore allocation and developmental observations of chimeras are not captured if social behaviour is defined based on spore bias alone, which led us to classify all evolved lines as cooperators. As already argued in Buttery et al (Buttery et al., 2009), Equation 2 is a good estimator of reproductive success only as long as strains produce, in isolation, comparable amounts of spores, which is not the case for the Ancestor and evolved lines (Figure S6). If one of the two populations that are mixed tended to form less spores, indeed, its expected proportion in the spores of a chimera would also be decreased. We can compute such expectation under the assumption that each strain contributes the same fraction of spores in isolation and in chimeras, hence the decision to turn into spores is independent of the presence of the other strain, as explained in Methods (Measures of spore formation efficiency and metrics of social performance).

Comparison of spore formation efficiency (percentage of cells that eventually become spores) to its expected value (Equation 4), computed based on that in clonal aggregates (Equation 1) allows us to distinguish the social behaviour between treatments (Figure 5 A). When the Ancestor strain is mixed with Bottom lines, the latter produce a smaller amount of spores than expected, while the opposite is true for the Ancestor strain. This scenario is consistent with the production of longer stalk, enriched in Bottom cells. In contrast, Top lines produce the same amount of spores as expected when alone (even more for the Top line 3, which was not able to produce spores on its own). Top lines however drastically reduce the amount of spores that the Ancestor is able to form, as they segregate out of the pre-stalk region, leaving the ancestor alone in providing the social function. Nonetheless, Ancestor cells become over-represented in the spore pool by the sheer fact that they are always part of the fruiting bodies - whereas Top cells get disproportionately excluded.

**Figure 5.**
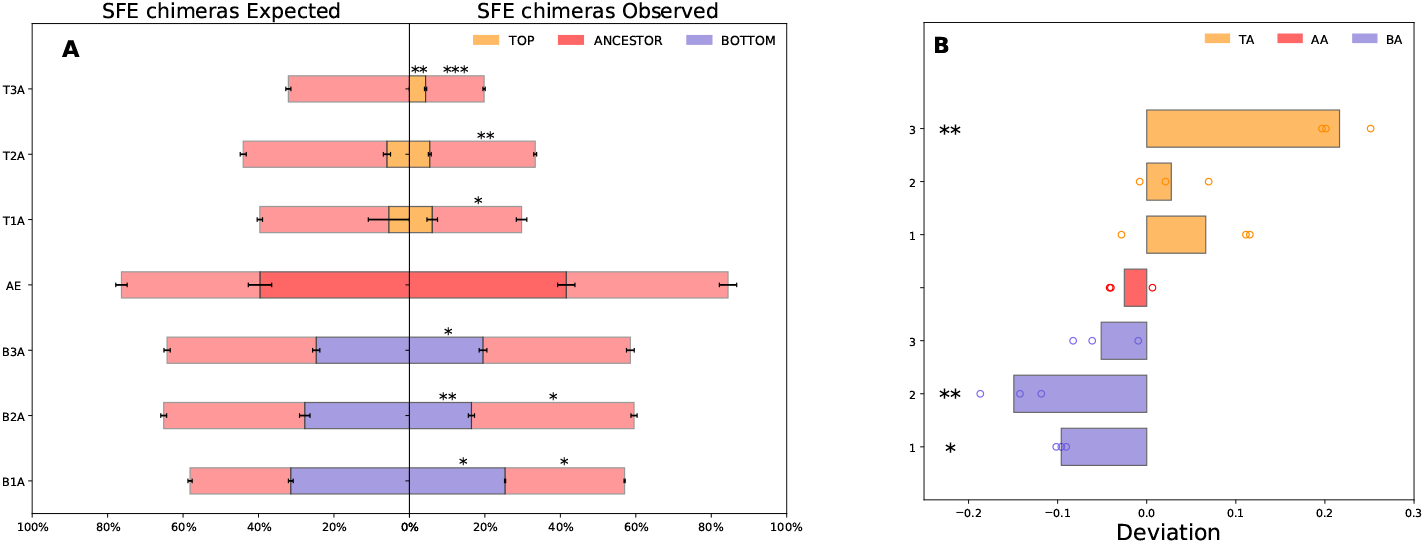
Spore formation efficiency in chimera and deviation: (**A**) Expected (left) and Observed (right) spore formation efficiency (SFE) in chimeras. Number of spores of the two co-aggregating strains relative to the number of cells of that strain in the mix (circles represent data points of three biological replicates and the RFP-tagged Ancestor strain is indicated in darkpink). Spore formation efficiency is significantly reduced when the Ancestor strain is mixed with both Top and Bottom lines (Student *t-test* comparison between Expected and Observed: * *p<*0.05, ** *p<*0.001, *** *p<*0.0001). (**B**) Deviation of spore allocation, computed according to Equation 4. Phenotypes of Top and Bottom lines are associated to opposite deviations of spore allocation: Top lines form more spores than expected whereas Bottom lines form less spores than expected. Although the trends seem clear-cut, deviations are however significant only for half of the evolved lines. However, the mean deviation of Top treatment is significant to the mean deviation of Bottom treatment and mean Ancestor deviation. (Student *t-test* mean significantly different to zero: * *p<*0.05, ** *p<*0.001). Colour code as in Figure 1.

These differences between Top and Bottom lines are captured when one compares the realized fraction of cells of a given type to its expected value, rather than to the fraction of this type in the initial mix. Equation 4 in the Methods (Measures of spore formation efficiency and metrics of social performance) generalizes Equation 2 when the expected probability of producing a spore differs between the two cell populations. Figure 5 B shows that this metric evidences that Top and Bottom cells differ in their social strategy. Indeed, while Bottom cells produce less spores than expected based on their own reproductive capacity, Top cells increase their proportion of spores with respect to the expectation. This is due to the fact that Top lines accomplish a close-to-normal development thanks to the Ancestor. Such exploitation is however not very efficient (and the advantage gained small) because, localizing in the pre-spore back region of the slug, Top cells expose themselves to segregation, which happens when the Top-rich tail of the slug brakes off and cannot undergo further differentiation. When comparing the mean deviation of each evolved line to zero, the differences are not all statistically significant. However, the mean deviation of the Top treatment was significantly different from both the Bottom treatment and the Ancestor. The qualitative pattern that emerges suggests that the use of other metrics for social success, along with the classical measure of spore bias, provides additional relevant information on the nature of the mechanistic processes underpinning social behaviour in cellular populations.

## DISCUSSION

In this work, we explored how selection acting on single-cell properties affects collective behaviour in cellular populations. Understanding how single-cell properties shape adaptations to collective living is key to explore possible evolutionary scenarios leading from unicellular to multicellular life cycles (Del Angel et al., 2020; Pentz et al., 2020; Márquez-Zacarías et al., 2021).

In our experimental design, selection on cell-substratum adhesion is applied to the ‘social amoeba’ *D. discoideum* at the unicellular stage of its life cycle, thus it does not act directly on social traits. Three independently evolved lines were obtained for each of two treatments, where cells were selected for higher (Bottom) and lower (Top) adhesion to the surface of a culture flask. Given the relatively small divergence time from their common ancestor and the small mutation rate (2,9.10^-11^ for the nuclear DNA (Saxer et al., 2012)), we expect that evolved lines and the Ancestor strain retain an almost complete genetic identity. Direct selection of a quantitative cell trait allowed us to factor out the effects of modified adhesion from those of broader genetic differences that occur in natural strains, and, unlike in mutagenized strains, to connect trait difference to their generative selection process. Chimeras containing the evolved lines and the Ancestor strain were used to assess the extent to which social behaviour can evolve as a byproduct of evolution of cell-level physical properties (Del Angel et al., 2020). We could interpret the resulting patterns of cell sorting based on associated variations in cell-substratum and cell-cell adhesion, and thus access the mechanisms underpinning alternative measures of social performance. In particular, we showed that different metrics provide a different classification of a strain’s behaviour.

Cellular adhesion has been frequently associated to ‘social behaviour’ in chimeras, as it can support assortment of cooperative cells. Different forms of cell-cell adhesion are involved in the assortative processes in the course of aggregation and development, and have potential evolutionary bearings. Mechanisms that allow genetically similar cells to recognize one another are often invoked in explaining how kin selection can maintain cooperative behaviour in spite of genetic heterogeneity. For instance, Tigr proteins provide a lock-and-key mechanism for specific recognition of cells that share the same allele at that locus (Hirose et al., 2011; Gruenheit et al., 2018). Similarly, the *csA* gene, that encodes a cell adhesion protein anchored in the cell membrane, has been proposed to act as a ‘green beard’ signal associated to cooperative behaviour (Queller et al., 2003). It has nonetheless been pointed out that non-specific adhesion can also play a constructive role in the evolution of social behaviour, and could have been determinant in early stages of the evolution of aggregative (as well as clonal) multicellularity (Ispolatov et al., 2012; Garcia et al., 2014, 2015; van Gestel and Wagner, 2021). Indeed, adhesion to surface is likely to be an ancestral feature that predated the emergence of multicellular organization. Moreover, cell sorting, a central process during development, is affected by cell-substratum adhesion through variations in interfacial tension, to which it is coupled through the acto-myosin cortex (Steinberg, 1975; Brodland and Chen, 2000; Brodland, 2002). However, the possible implication of adhesion to surfaces in the evolution of aggregative multicellularity has not, to our knowledge, been previously explored in *D. discoideum*.

A first form of social behaviour manifests in the multicellular cycle of cells of the same genotype. Modifications in cell-substratum adhesion produced by knocking out specific genes was reported to affect aggregation, developmental timing and differentiation (Plak et al., 2016). Similar effects of surface adhesion on developmental patterns were observed in our evolved lines, both in clonal development and in chimeras. Cells of the Bottom lines, more adhesive to the surface, were particularly enriched in the anterior of chimeric slugs. This is consistent with the previous observation that anterior cells display, upon disaggregation of the slug, stronger adhesion than posterior cells (Yabuno, 1971), suggesting that cell sorting in the slug is generally influenced, directly or indirectly (for instance, through a correlated change in polar cell adhesion) by cell-substratum adhesion systems. The Top lines, less adhesive to the surface, differed from the Ancestor in other ways. During clonal aggregation, they displayed patterns similar to those obtained when cell-substratum adhesion was reduced by altering the chemical properties of the surface (Wang et al., 2014), and the timing of clonal development was severely altered. In chimeras, Top cells distributed in the slug like strains where adhesion to the surface was impaired by knocking out the *paxB* or *dimA-* genes, which impact cell sorting or cell fate, respectively (Bukahrova et al., 2005; Foster et al., 2004).

Spore production in chimeras of strains with different genetic background – typically natural isolates or mutagenized variants – mixed pairwise are the cornerstone of experimental use of *Dictyostelium* and, more generally, of cooperation in microbes, to test evolutionary theories. Given the ease of counting spores that derive from one or another strain, social behaviour has been extensively associated to the over-representation (‘cheating’) or under-representation (‘cooperation’) of one genotype in the spore pool (Fortunato et al., 2003a; Foster et al., 2004; Santorelli et al., 2008, 2013). We have shown that cooperative variants are promptly evolved when selection acts on single-cell adhesion, suggesting that in natural populations evolution against cooperation could be compensated by evolutionary processes acting on single cells.

The action of selection on properties of isolated cells has been so far largely overlooked when considering the evolution of social traits, with the notable exception of the role of ‘loner’ cells (Dubravcic et al., 2014; Rossine et al., 2020; Miele and De Monte, 2021). Here, we showed that properties of isolated cells can be important not only in fluctuating environments, where different behavioural patterns are effective in hedging cells’ bets in front of unpredictable selective pressures. Through their direct and indirect consequences on the multicellular phase of the life cycle, changes in cellular physical properties can oppose selection on multicellular traits and constrain the possible solutions available to development.

Our observations revealed that the definition of social behaviour based purely on the fraction of spores produced can actually conceal divergent developmental paths. When spore bias is computed as the distance of spore frequency with respect to cell frequency in the mix (Equation 2) both Bottom and Top lines come to be classified as cooperators. However, observation of the developmental cycle revealed fundamental differences in contribution of the evolved lines to multicellular function. Bottom cells appear to ‘sacrifice themselves’ by forming the upper cup and the basal disk, which are both essential for the formation and the structural stability of the fruiting body. Cooperative behaviour in this case can thus result from stronger attachment to the surface and increased polar contacts. Top cells, on the other hand, appear to act within the slug like cheaters, as they tend to be found in the back of the slug and lead to a decrease in the total spore production of the partner. This strategy can be fruitful as long as Ancestor cells can compensate for the presence of a small proportion of Top cells. Further studies are underway to check if, along the lineage of the Top treatment, lines that evolved only a slight decrease in adhesion produce a spore bias compatible with unconditional cheating. In the conditions we have examined, lack of coordination between cells results in a counter-productive detachment of Top-enriched tails of streams and slugs. In this way, Top cells end up undermining not only the reproductive success of the Ancestor, but their own as well. Recognition that apparent cooperation can come as a side effect of excessive greed is captured by metrics that, in computing the expected composition of the spore pool, also consider the amount of spores produced by each strain on its own (Equation 4). This metric (as that proposed in Buttery et al. (Buttery et al., 2009)) takes into account that one strain may produce more spores in the first place, which would give it a head start in the chimera if the decision of a cell to turn into a spore was independent of the social interactions. The second definition seems to better reflect the developmental differences observed in the evolved lines.

During the vegetative stage, amoebas of *Dictyostelium* crawl on surfaces as single cells. Several genes have been found to play a role in cell–substratum adhesion, among which some *TalB, PaxB* knock out mutants are implicated in development and cell sorting (Tsujioka et al., 1999; Bukahrova et al., 2005; Cornillon et al., 2008, 2000) (for a review of current knowledge about cell-substratum adhesion in *Dictyostelium discoideum*, see (Mijanović and Weber, 2022)). However, their interactions with cell-cell adhesion, whose role is limited to multicellular development, are not yet characterized. In our work, we have identified putative pleiotropic effects of changes in cell-surface adhesion on cell-cell adhesion. Even though we do not know what are the molecular basis of such co-variation, we showed that it is sufficient to qualitatively explain the developmental patterns and social behaviour observed in chimeras. Pleiotropy is a recognized mechanism acting within a single life cycle to counterbalance selective pressures acting on specific phenotypic traits. It has been for instance invoked as the reason why mutants with decreased sensitivity to DIF (differentiation-inducing factor), a morphogen controlling the stalk-specific pathway of differentiation, are not positively selected (Foster et al., 2004; Strassmann and Queller, 2011). Despite being considered an important feature of systems where cooperation is mediated by public goods (Dandekar et al., 2012), the way pleiotropic effects deploy along an evolutionary trajectory is seldom addressed. Our experiment allows to reveal how pleiotropic effects can steer social interactions in unexpected directions.

Evolutionary effects similar to those we evidenced could be produced by a number of other selective pressures that, acting outside the multicellular phase of the life cycle, end up affecting the way cells coordinate spatially and temporally to achieve a collective function. Possible future investigation may consider whether cell-intrinsic differences in motility or response to the environment are as effective as adhesion in affecting the course of social evolution.

## Materials and methodss

### Strains and culture condition

*Dictyostelium discoideum* axenic AX3 strain (Dictybase ID: DBS0235545) was transformed with plasmids pTX-GFP (Dictybase ID: 11) or pTX-RFP (Dictybase ID: 112) to express either GFP or RFP fluorescent markers respectively. The fluorescent proteins encoded on the plasmid also carries a gene for antibiotic resistance (Gentamicin 418, Sigma-Aldrich: G418). Cells were cultured in autoclaved HL5 medium (per L, 35.5 g HL5 from formedium, pH=6.7) at 22°C and with a concentration of 20 μg mL^−1^ G418.

### AX3 strain transformation

GFP and RFP-expressing cell strains were obtained by transforming cells as in Dubravcic et al (Dubravcic et al., 2014). Briefly, cells were transformed using a standard electroporation procedure with pTX-GFP or pTX-RFP. AX3 cells were grown in 75 *cm*^2^ flasks until they reach high density (but before they become confluent). Four to six hours before the transformation, the medium was changed. Cells were then re-suspended in 10 mL of ice-cold HL5 and kept on ice for 30 minutes. Cells were centrifuged for 5 minutes at 500 g and 4°C. The pellet was re-suspended in 800 μL of electroporation buffer and transferred into ice cold 4 mm electroporation cuvettes containing 30 μg of plasmid DNA. Cells were electroporated at 0.85 kV and 25 mF twice, waiting for 5 seconds between pulses and transferred from the cuvette to 75 *cm*^2^ flask with HL5 medium. To select transformants, the next day 5 μg mL^−1^ of the antibiotic G418 was added to the culture media. The concentration of G418 was gradually increased from 5 μg mL^−1^ to 20 μg mL^−1^ over 1–2 weeks and resistant cells were collected and frozen.

### Evolution of cell-substratum adhesion

Evolution experiments were carried out for two months by performing thirty rounds of selection by serial transfer of cells contained either in the supernatant (Top) or sticking to the culture vial after removal of the liquid (Bottom) (Figure 6). Starting from a common ancestor, three replicate lines were independently subjected to the Top and Bottom selection treatments.

**Figure 6.**
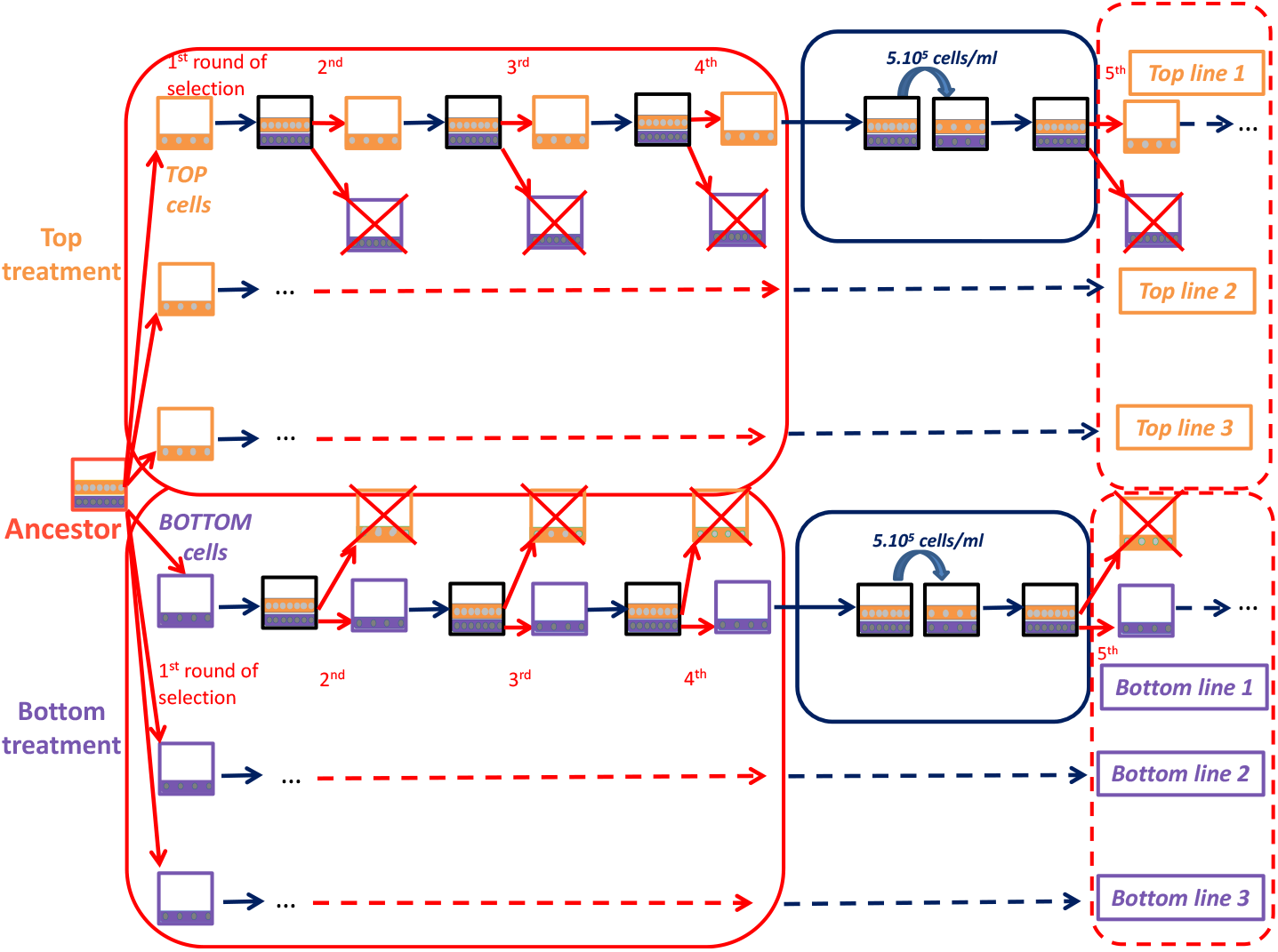
Schematic representation of the Evolution experiment. Artificial selection was applied to the cell-substratum adhesion phenotype, according to two distinct treatments. Top lines were obtained by serial transfer of cells in the supernatant (orange). Bottom lines were obtained by serial transfer of cells attached to the bottom flask (violet). The complement was discarded at each selection step (red arrows), and the selected component was diluted in fresh medium and let to grow for a day (blue arrows). Cells were subjected to two transfer regimes: selection was applied once per day for four consecutive days (red boxes: four rounds of selection); cells were then let grow freely for two days (blue box), so as to maintain a healthy and growing population of cells.

Cells were initially inoculated in 10 mL of fresh HL5 medium with 20 μg mL^−1^ G418 in 25 *cm*^2^ flasks at a density of 5.10^5^ cells/mL. After one day of growth, the Top (Bottom) components were separated to produce a fresh, selected cell population as follows: (i) In the Top treatment, the culture flask was lightly shaken to detach cells which were not strongly attached to the bottom surface and to homogenize. A volume of 5 mL of the liquid was then transferred in 5 ml HL5 with 20 μg mL^−1^ G418 in 25 *cm*^2^ flasks. (ii) In the Bottom treatment, cells that remained attached to the culture flask after lightly shaking the culture flask and removing the liquid were removed by washing the flask with 10 mL HL5 with 20 μg mL^−1^ G418. Cells were detached by 10 to 15 cycles of pipetting with a Pasteur Pipette. A volume of 5 mL of the resulting cell suspension was then transferred in 5 mL HL5 with 20 μg mL^−1^ G418 in 25 *cm*^2^ flasks.

For both treatments, replenished cultures were let grow for a day before another round of selection. This selection was applied for four consecutive days, after which cells were let grow without selection for two days. Before resuming the selection protocol, the whole culture was diluted to a density of 5.10^5^ cells/ml. The thirty rounds of selection correspond roughly to 85 and 97 generations (estimated in Figure S1) for the Top and Bottom treatment, respectively. During this experiment, cells were kept in vegetative growth, so that selection only acted on single-cell, and not on multicellular traits.

### Cell-substratum adhesion assay

Efficiency of cell-substratum adhesion was estimated by measuring the fraction of cells attached to a culture flask. If *N* the total number of cells in a culture and *n*_*L*_ those that are suspended in the liquid, this efficiency is quantified as (*N* − *n*_*L*_)*/N*. Such quantity was measured (see protocol below) in three different conditions: 1) at the beginning of exponential growth (24 hours after initiating a culture at 5.10^5^ cells.mL^−1^); 2) right after starvation, where cells at the beginning of exponential growth were washed out of the nutrient medium by three successive centrifugations with SorC buffer (per L, 0.0555g CaCl_2_; 0.55g Na_2_HPO_4_,7H_2_O; 2g KH_2_PO_4_) at 500 g for 7 min; 3) during streaming, where cells were collected at the streaming stage of the developmental cycle (whose timing was identified by analysing time-lapse movies) by washing plates with 1 mL SorC to disaggregate clusters. Cultures in these three conditions were re-suspended to a density of 5.10^5^ cells.mL^−1^ in 10 mL medium (HL5 with 20 μg mL^−1^ G418 for condition 1 and SorC for conditions 2 and 3) in 25 *cm*^2^ flasks. This concentration was low enough so that cells could independently attach to the surface. The total cell number *N* was then counted using a hemocytometer. Cultures were let rest for 30 minutes, after which flasks were lightly shaken to detach cells that were not strongly attached to the bottom. The supernatant was collected into 15 mL tube and the number of cells in suspension *n*_*L*_ measured with a hemocytometer.

Cell-substratum adhesion was measured in condition 1) for the three evolved Top and Bottom lines and for the Ancestor. In order to estimate variations of cell-substratum adhesion during development (conditions 2) and 3)), these measures were performed for line 1 of the Top and Bottom treatments and the Ancestor. All measures have been realized in triplicates, starting from the same culture.

### Cell-cell adhesion assay

Cell-cell adhesion was quantified by the method of Gerisch (Gerisch, 1968). The principle of this method is to allow cells to get in contact with each other in a shaken culture – thus independent of their substratum adhesion –, and then measure the proportion of cells that are found in clusters. The percentage of clustered cells was determined by subtraction from the total cell number of the number of unaggregated cells, and then dividing by the total cell number. The cell culture was centrifuged and re-suspended at a density of 2.10^6^ cells.mL^−1^ in 2 mL of SorC buffer. This cell density was chosen so that only a fraction of cells aggregate. We explored the mechanisms responsible for changes in cell adhesion observed in this work by reasoning that changes in cell-cell adhesion are unlikely to involve newly evolved contact sites, and that selection has probably acted on existing adhesion systems.

Quantification of the fraction of aggregated cells was previously used to address the role of two families of proteins that affect cell-cell adhesion during *Dictyostelium* development. The adhesion system DdCAD-1 (gp24) is expressed soon after starvation and is inhibited by Ethylene Diamine Tetra acetic Acid (EDTA) (Sriskanthadevan et al., 2011; Hou, 2004). Later, in the streaming and mound phases, polar cell adhesion by EDTA-resistant gp80 glycoproteins causes cells to organize in chains (Fontana, 1993) and elicits activation of the cAMP-mediated signal transduction pathway (Desbarats et al., 1994).

Addition of EDTA to the culture, together with the observation in two different developmental phases, informed us on the part that mechanisms involved in the early stages of multicellular development have on changes in cell-cell adhesion of derived lines.

In the EDTA assays, 10 mM EDTA were added. The culture was rotated at 150 rpm and 22°C during 45 minutes. The total number of cells and number of unaggregated cells were subsequently measured using a hemocytometer. The counting protocol was repeated for cultures prepared in conditions 2 − 3 as described in ‘Cell-substratum adhesion assay’, and in triplicate for the Ancestor and each of the three Top and Bottom lines.

### Developmental life cycle

Cells were grown in 25 *cm*^2^ flasks with 10 mL HL5 medium with 20 μg mL^−1^ G418 until they reached the mid-exponential growth phase. Cell density at the onset of starvation was comparable for all starvation assays. Cells were washed from the nutrient medium by three successive centrifugations with SorC buffer at 500 g for 7 min. After the last centrifugation, the pellet was re-suspended in SorC buffer and the concentration adjusted to 2.10^7^ cells.mL^−1^. A volume of 40 μL (corresponding to 4.10^6^ cells/cm^2^) of suspension was plated on 6 cm plates filled with 2 mL of 2% Phytagel (Sigma-Aldrich) as previously described by Dubravcic et al. (Dubravcic et al., 2014). We produced chimeras by mixing equal amounts of each evolved line (marked with GFP) with the Ancestor strain (marked with RFP). The concentration was adjusted to 2.10^7^ cells.mL^−1^ and a volume of 20 μL of each was mixed before plating (corresponding to a total of 4.10^6^ cells/cm^2^ plated). The developmental cycle was subsequently imaged as explained in paragraph ‘Time-lapse microscopy and image analysis’. We first checked that the different fluorescent markers did not give rise to segregation during aggregation or development (Figure 4 middle (AA); Movie S11: multicellular developmental cycle in a chimera between Ancestor GFP and RFP). Consistent with the observed lack of bias in the spores, no distinctive pattern was recognizable at any developmental stage and the developmental timing was comparable to that of single Ancestor populations (Figure S10).

### Time-lapse microscopy and image analysis

Cells were starved as described in section ‘Developmental life cycle’. The 6 cm diameter Petri dish was imaged on an automated inverted microscope Zeiss Axio Observer Z1 with a Camera Orca Flash 4.0 LT Hamamatsu and using a 5X objective. Images were acquired with MicroManager 1.4 software. This setup allows Petri dish scanning at regular time intervals (typically 5 min), with phase contrast and fluorescence image acquisition (with 33 ms exposure times). A mosaic image is reconstructed by combining all the images of contiguous areas of the Petri dish at a given time point using a custom Python program. Time-lapse movies were captured in duplicates for clonal (Top, Bottom, Ancestor) and chimeric development (Top-Ancestor, Bottom-Ancestor, Ancestor GFP - Ancestor RFP) for each evolved line.

#### Timing of development

By visually inspecting the time-lapse movies for Top, Bottom and Ancestor lines and their chimeras, six characteristic developmental stages were identified : streaming, mound, tipped aggregate, slug, Mexican hat and fruiting body. The moment at which each developmental stage appeared for the first time was used to partition the developmental cycle.

#### Aggregate size and number

Aggregates size and number change during the aggregation process. In order to compare different movies, a custom imageJ macro (Schindelin et al., 2012) was written to compute them at the time point when their values stabilize, indicating the end of the aggregation process. At this point, aggregates area was averaged over the whole aggregation domain.

#### Cell position in binary chimeras during developmental life cycle

Time-lapse movies were realized as previously described also for 1:1 binary chimeras where either GFP-labelled evolved lines or the GFP-labelled Ancestor (for the control) were mixed with RFP-labelled Ancestor cells.

### Measures of spore formation efficiency and metrics of social performance

Evolved lines (three Top and three Bottom) and Ancestor strain were subjected to the developmental life cycle either clonally or in binary chimeras (as described in section ‘Developmental life cycle’). In addition, ancestral RFP and ancestral GFP were mixed to control for the possible differential effects of different plasmid insertion. Each experiment (clonal Ancestor and evolved lines, and their chimeras with the Ancestor) was realised in triplicate.

The initial number of cells plated was estimated using a hemocytometer. At the end of the developmental life cycle, spores were collected using 1 mL pipette tips after washing the plates with 1 mL SorC. The number of spores was counted using the hemocytometer. Spore Formation Efficiency *SFE*_*i*_ of a strain *i* is measured as the ratio between the number *N*_*i*_ of spores produced by that strain and the number of cells of the same strain plated at the beginning of the experiment *n*_*i*_:

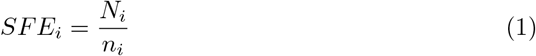

Here, lower and upper case indicate the composition at the beginning (*n*) and at the end (*N*) of multicellular development, the latter being the moment when is established the spore pool that will seed the following generation.

In chimeras, the proportion of cells of the two types was quantified using a flow cytometer Cube8 Partec with SSC (Side Scatter), FSC (Forward Scatter), FL1 (GFP) and FL3 (RFP) channels. The proportion in the plated population was assessed by diluting 10 μL of the initial suspension in 990 μL of SorC. The proportion in the spores was similarly measured. Since the cytometer does not clearly differentiate between spores and non aggregated cells, the latter were eliminated by adding first 0.05% SDS for 5 minutes followed by a centrifugation at 250 g for 3 minutes. The pellet was re-suspended in 1 mL SorC before passage in the cytometer.

For binary mixtures with strain *j* (written as a superscript in formulas), social performance of strain *i* (written as a subscript in formulas) was quantified in two ways, starting from the measure of the fraction of spores 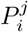:

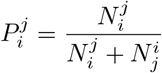

and the fraction 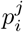 of plated cells of strain *i* when in chimera with the strain *j* :

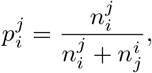

that was close to 50%. First, following classical estimation of social behaviour (Fortunato et al., 2003a; Foster et al., 2004; Santorelli et al., 2008, 2013), we simply assessed the deviation in strain’s frequency during multicellular development (Spore Bias: ‘SB’):

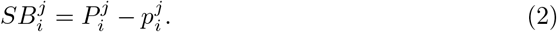

This metrics is commonly used as a measure of relative fitness of strain *i*, as it quantifies the difference between the probability that a cell of type *i* becomes a spore with respect to the frequency of that type in the population. However, as noticed by Buttery et al. (Buttery et al., 2009), this formula is a good estimator of reproductive success only as long as strains produce in isolation a comparable amount of spores.

Indeed, if strains can be found in clonal as well as chimeric groups, the social performance should also take into account the spore production success of strains in isolation. In order to take into account differences in spore production between strains, Equation 2 can be generalized by estimating the number of spores of type *i* that are expected, based on each strain’s SFE in isolation (Equation 1), assuming that the probability that a cell of one strain turns into spores is the same in chimeras as for clonal aggregations. The expected fraction of spores in chimeras can thus be estimated based on the spore production in clonal development as follows:

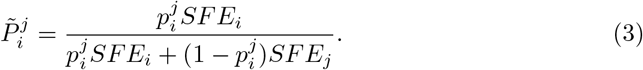

The expected proportion of spores thus depends on the initial frequency of the two strains.

Social performance of strain *i* in a chimera with strain *j* is then defined as the deviation of the observed proportion of *i* spores with respect to its expected value:

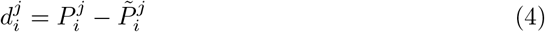

Notice that if the two strains have the same spore formation efficiency, then the two metrics are equivalent: 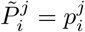 and 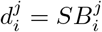. Moreover, it is easy to see that in binary mixes 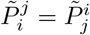, so that this metric respects the symmetry expected for relative fitness advantages.

For both metrics, a negative value indicates a decreasing contribution to the spore pool and is considered as indication of a strain’s cooperative behaviour. On the opposite, a positive value indicates a higher contribution to the spore pool than expected, which is indicative of social cheating.

### Statistical analysis

Statistical analysis was performed in R Studio. Significance of pairwise comparison between the samples was established using two-sample Student *t-test* function, and a *p*-value *p<*0.05 was considered as significant.

## Author contributions

Conceptualization, Methodology, Formal analysis: S.A; Investigation: S.A and M.F; Writing – Original Draft and revised manuscript: S.A and S.D.M.

## Acknowledgments

The authors are very grateful to Clement Nizak for helpful discussions. This work has received support under the program “Investissements d’Avenir” launched by the French Government and implemented by ANR with the references ANR–10–LABX–54 MEMOLIFE and ANR–10–IDEX–0001–02 PSL* Université Paris, Q-life ANR-17-CONV-6150005, and the project ANR-19-CE45-0002 ‘ADHeC’ PSL research University.

## Declaration of Interests

The authors declare no competing interests.

## Notes

### Competing Interest Statement

The authors have declared no competing interest.

### Summary of Updates

Data with three evolved lines

