## Supplementary Information for "Evolving social behaviour through selection of single-cell adhesion in *Dictyostelium discoideum*"

### Supporting information

*Figure S1.* **Number of generations of Ancestor strain and Top and Bottom lines 1.**

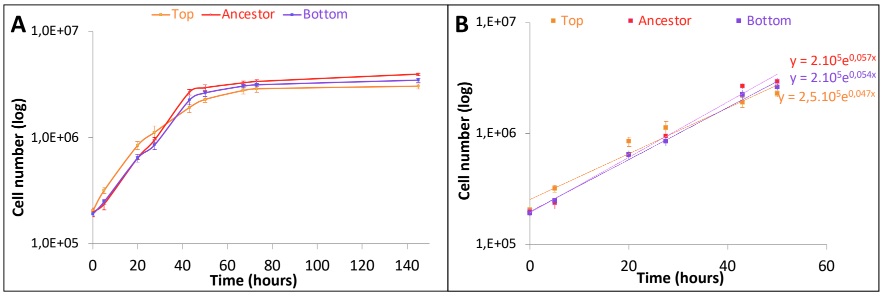


The doubling time $DT$ was calculated at the end of the evolution experiment for Top and Bottom line 1 by estimating the exponential growth rate. The growth curve (panel A, number of cells measures with the hemocytometer) indicates that cells grow approximately exponentially for 50 hours, before saturation of the population size. This results in linear growth in a semi-logarithmic plot (panel B), the slope $a$ of which was computed by linear regression ($a$= 0.047; 0.054 and 0.057 for Top, Bottom and Ancestor respectively). From this, we estimate the doubling time $DT = \frac{ln2}{a}$) as 14.6 hours, 12.8 hours and 12.1 hours for Top, Bottom and Ancestor respectively. So, selection had an effect on growth rate, but changes were likely contained by the experimental design, that alternated selective passaging with undisturbed growth. Overall, the difference in number of generations was modest. The evolution experiment, indeed, was performed during $D = 52$ days, corresponding to a number of generations ($G = \frac{D}{DT}$) approximately equal to 85 and 97 for Top and Bottom, respectively. (colour code as in Figure 1).

*Figure S2.* **Spore formation efficiency of chimeras composed of evolved and Ancestor strains.**

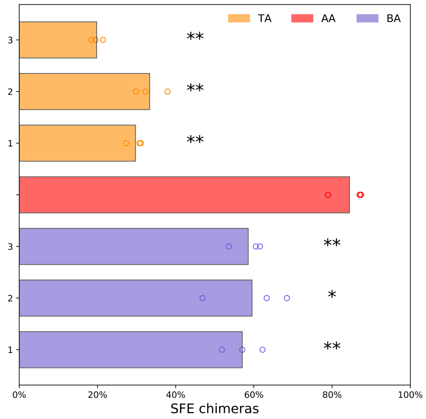


Chimeras composed of Top (TA) and Bottom lines (BA) mixed with the Ancestor formed less spores compared to chimera composed of Ancestor strains (RFP and GFP, (AA)). (colour code as in Figure 1. Circles represent data point of three biological replicates. Student *t-test* significant compared to the chimera AA: * $p$<0.05; ** $p$<0.001).

*Movie S3: Clonal aggregation of the Ancestor, Top (line 1) and Bottom (line 1).* Phase contrast images: Developmental life cycle of Ancestor (left) , Top (middle) and Bottom (right). The beginning of the different stages of the multicellular developmental cycle (summarized in Figure 2) is indicated.

*Figure S4.* **Clonal and chimeric development of evolved lines 2 and 3.**

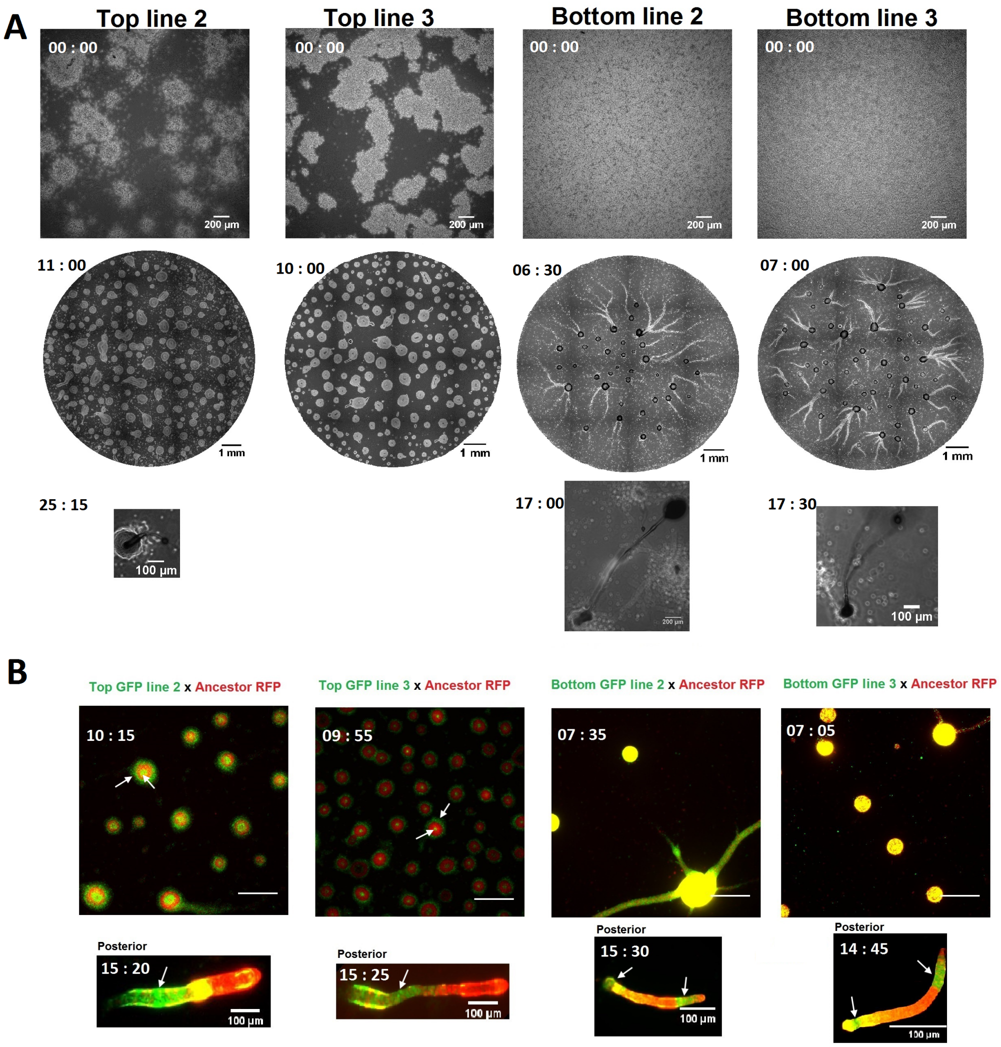


**(A)** Phase contrast images of three stages of the cycle: starting aggregation, mounds, fruiting bodies. Pictures represent Lines 2 and 3 of the Top and Bottom treatments. **(B)** Developmental patterns in chimeras (aggregation and slug stage) for Lines 2 and 3 of the Top and Bottom treatments (green) when mixed with Ancestor RFP (red). (Scale bar = 500 µm)

Differences in clonal and chimeric development are discussed in the text taking the Line 1 as representative for each treatment.

*Figure S5.* **Mean area ($mm^2$) and number of aggregates in clonal aggregations.**

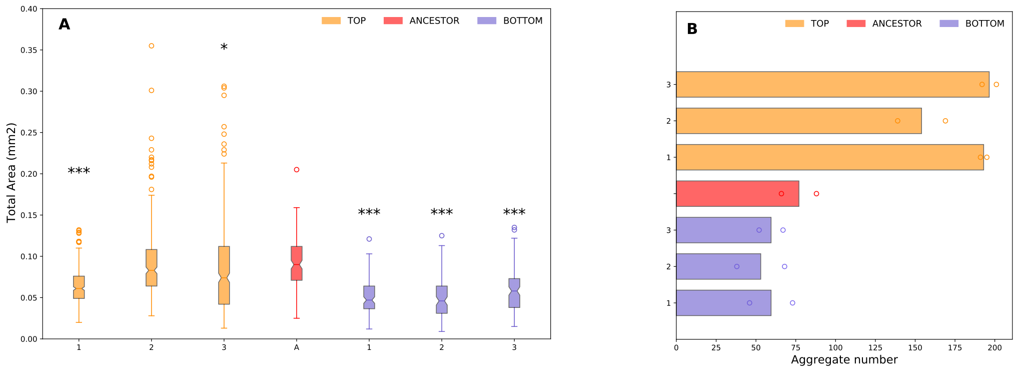


**(A)** Mean aggregate area (averaged over two independent replicates). Aggregates were smaller for Top and Bottom compared to Ancestor (Student *t-test* significant compared to the mean of the Ancestor: * $p$<0.05, ** $p$<0.001, *** $p$<0.0001). **(B)** Number of aggregates was higher for the Top treatment compared to Bottom treatment (Student *t-test* $p = 3.10^{-6}$). Circles represent data points of two biological replicates.. Colour code as in Figure 1.

*Figure S6.* **Spore formation efficiency in evolved lines and Ancestral strains.**

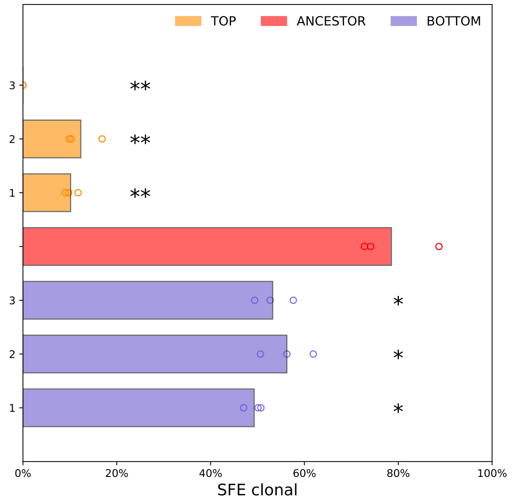


As a result of impaired development, both Top and Bottom evolved lines produce fewer spores than the Ancestor. (Student *t-test* significant compared to the mean of the Ancestor: * $p$<0.05, ** $p$<0.001, *** $p$<0.0001, colour code as in Figure 1). Circles represent data points of three biological replicates.

*Figure S7.* **Cell-substratum adhesion during growth, at starvation and during streaming for Top and Bottom lines 1 and Ancestor.**

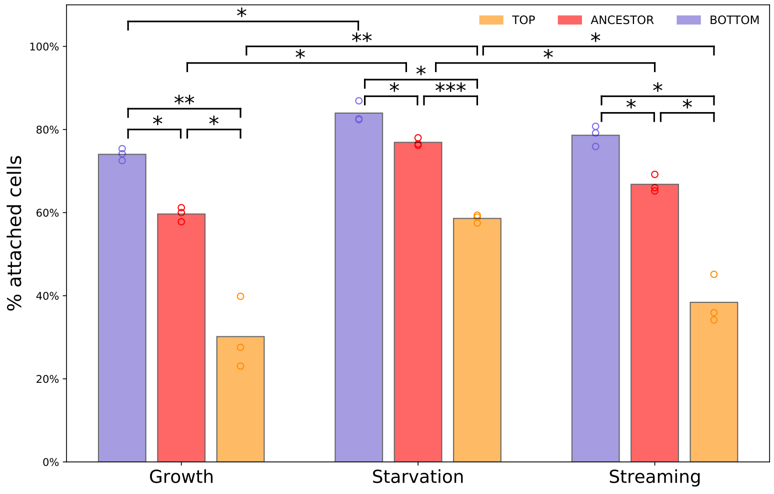


We measured cell-substratum adhesion during exponential growth (when the selection was performed) and at two additional time points. 'Starvation' is just after cells were transferred into buffer (after centrifugation), before being plated to start the developmental cycle, and is the same for all lines. The onset of the streaming phase, instead, corresponds to different times after plating, depending on the treatment: five hours for Ancestor and Bottom cells and seven hours and a half for Top cells. Measures at such 'Streaming' time point were realized on cells collected from a plate, where they had started their developmental cycle. Cell-substratum adhesion differences remain qualitatively the same at different time points: Top cells remain the less adhesive and Bottom cells the most adhesive to the surface. In accordance with previous studies (Rosengarten et al., 2015), cell adhesion is however modulated along the developmental life cycle. With respect to exponential phase in rich medium, cell-substratum adhesion is increased after starvation in all treatments, and decreases (in a more pronounced way in Top cells) at the streaming stage. Although cell-substratum adhesion changes in time, qualitative differences among strains are maintained. (Circles represent data points of three biological replicates. Colour code as in Figure 1; Student *t-test*: * $p$<0.05, ** $p$<0.001, *** $p$<0.0001).

*Figure S8.* **Cell-cell adhesion differences through development.**

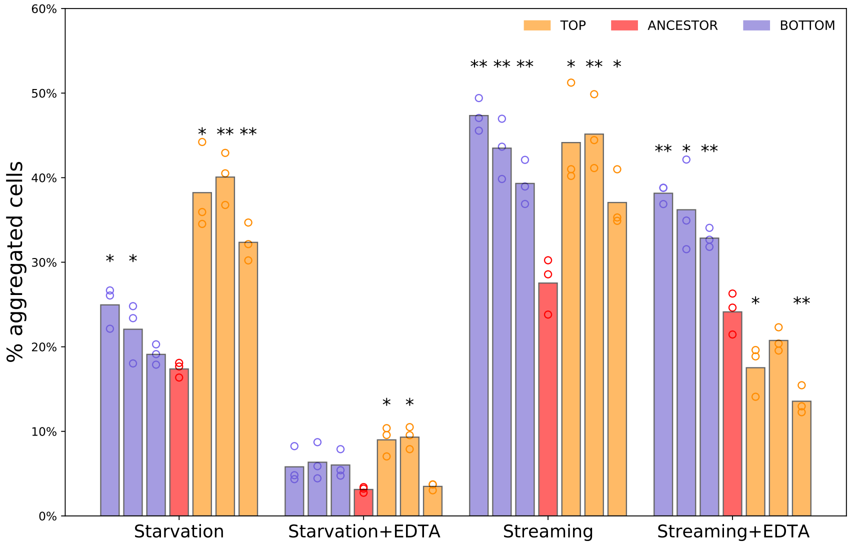


The percentage of aggregated cells at the beginning of starvation and during streaming is higher for the derived lines than the Ancestor. At starvation, EDTA drastically reduced cell clustering, as expected , in similar proportions for all lines. During streaming, EDTA produces a small decrease in clustering for Ancestor and Botttom lines, that seem thus to maintain their polar adhesion intact (but stronger for the Bottom lines). Instead, it is highly disruptive for the Top lines (most of all for line 3). This also explains why Top cells are more adhesive than Bottom earlier, while the reverse is true later. Differential decrease of such percentage upon addition of EDTA suggests that higher cell-cell adhesion of Top cells is mostly due to early-development adhesion proteins. It is consistent with the previous observations that Bottom cells form longer streams. Top cells' development, on the other hand, resembles that of mutants lacking the EDTA-sensitive gp80, characterized by breaking streams and smaller aggregates (Roisin-Bouffay et al., 2000). This suggests that the observed variation in cell-cell adhesion might be due to a lower expression level of the gp80 adhesion protein. Circles represent data point of three biological replicates. (Student *t-test* significant compared to the mean of the Ancestor: * $p<0.05$, ** $p<0.001$, colour code as in Figure 1).

*Table S9.* **Cell-cell adhesion: comparison between treatments without and with EDTA.** Student *t-test* *p*-value: comparison between starvation versus starvation with EDTA and streaming versus streaming with EDTA for each evolved line.

| | T1A | T2A | T3A | AA | B1A | B2A | B3A |
|---|---|---|---|---|---|---|---|
| Starvation | **0.006** | **0.00097** | **0.0015** | **0.0004** | **0.0007** | **0.0054** | **0.0006** |
| Streaming | **0.0074** | **0.0064** | **0.0019** | 0.233 | **0.0047** | 0.1344 | **0.0352** |

*Figure S10.* **Timing of the developmental life cycle in chimeras.**

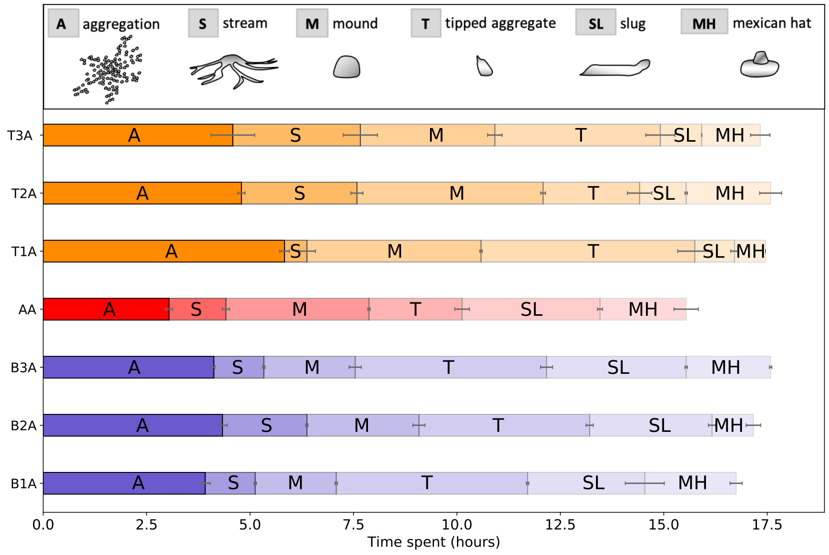


Chimeras of Bottom-Ancestor (BA); Ancestor GFP -Ancestor RFP (AA) and Top-Ancestor (TA), evolved lines 1, 2, 3. The three independently evolved lines from the same treatment behave similarly. AA behaves like the pure Ancestor. Each phase starts with the first apparition of the different multicellular stages, indicated on top in time-lapse movies (Movie S12: multicellular developmental cycle in a chimera between Top (line 1) and Ancestor, Movie S13: multicellular developmental cycle in a chimera between Bottom (line 1) and Ancestor). Colour code as in Figure 1.

*Movie S11: multicellular developmental cycle in a chimera between Ancestor GFP and RFP.* Time-lapse movies in phase contrast (upper left), GFP (lower left) and RFP (lower right) channels, and merged (upper right) during the multicellular life cycle.

*Movie S12: multicellular developmental cycle in a chimera between Top (line 1) and Ancestor.* Time-lapse movies in phase contrast (upper left), GFP (lower left) and RFP (lower right) channels, and merged (upper right) during the multicellular life cycle.

*Movie S13: multicellular developmental cycle in a chimera between Bottom (line 1) and Ancestor.* Time-lapse movies in phase contrast (upper left), GFP (lower left) and RFP (lower right) channels, and merged (upper right) during the multicellular life cycle.
